# Genetic loss of norepinephrine does not alter adult hippocampal neurogenesis in dopamine β-hydroxylase deficient mice

**DOI:** 10.1101/2022.09.14.508040

**Authors:** Darshana Kapri, Krishna C. Vadodaria, Karen S. Rommelfanger, Yvonne E. Ogbonmwan, L. Cameron Liles, Kimberly A. Fernandes-Thomas, Sonali S. Salvi, Basma F.A. Husain, David Weinshenker, Vidita A. Vaidya

**Author notes:** Equal senior corresponding authors Address correspondence to: Dr. Vidita A. Vaidya, Department of Biological Sciences, Tata Institute of Fundamental Research, Homi Bhabha Road, Mumbai 400005, India., Dr. David Weinshenker, Department of Human Genetics, Emory University School of Medicine, 615 Michael St., Whitehead 301, Atlanta, Georgia 30322, USA.

## Abstract

Norepinephrine (NE), and specific adrenoceptors, have been reported to influence distinct aspects of adult hippocampal neurogenesis, including latent stem cell activation, progenitor proliferation, and differentiation. These findings are predominantly based on the use of pharmacological approaches in both *in vitro* and *in vivo* systems. Here, we sought to assess the consequences of genetic ablation of NE on adult hippocampal neurogenesis, by examining dopamine β hydroxylase knockout (*Dbh-/-*) mice, which lack NE from birth. We find that *Dbh-/-* mice exhibit no difference in adult hippocampal progenitor proliferation and survival. Further, the number of immature newborn neurons, labeled using stage-specific developmental markers within the hippocampal neurogenic niche, was also unaltered in *Dbh -/-* mice. In contrast, the noradrenergic neurotoxin DSP-4, which had previously been shown to reduce adult hippocampal neurogenesis in rats, also resulted in a decline in hippocampal progenitor proliferation in C57/Bl6N mice. These findings indicate that pharmacological lesioning of noradrenergic afferents in adulthood, but not the complete genetic loss of NE from birth, impairs adult hippocampal neurogenesis in mice.

## 1. Introduction

Select neurogenic niches within the adult mammalian brain retain the ability to generate newborn neurons throughout life, including the subventricular zone (SVZ) lining the lateral ventricles and the subgranular zone (SGZ) in the hippocampal dentate gyrus (DG) subfield (Bond et al., 2015). Adult hippocampal neurogenesis encompasses the activation of quiescent neural progenitors, proliferation, fate specification, post-mitotic survival, migration, maturation, and functional integration into existing neurocircuitry (Bond et al., 2015). This multi-stage process is regulated by diverse intrinsic factors including growth factors, developmental morphogens, transcription factors, neurotransmitters, and hormones, as well as being modulated by extrinsic factors such as psychotropic drugs, exercise, and enriched environment, several of which are known to influence noradrenergic signaling (Bond et al., 2015). Previous work has identified norepinephrine (NE) as a key regulator of quiescent stem cell activation and progenitor proliferation in the hippocampal neurogenic niche (Kulkarni et al., 2002; Jhaveri et al., 2010; Coradazzi et al., 2016). Pharmacological lesion of noradrenergic afferents using the locus coeruleus (LC)-specific neurotoxin DSP-4 robustly reduces hippocampal progenitor proliferation in adult rats (Kulkarni et al., 2002; Coradazzi et al., 2016). Further, *in vitro* studies have shown that NE directly stimulates a multipotent and self-renewing neural hippocampal progenitor population, and that antidepressants that target the NE transporter exert significant neurogenic effects (Jhaveri et al., 2010). Adrenergic receptors (ARs) are expressed by hippocampal progenitors within the neurogenic niche, and β3AR activation enhances hippocampal progenitor turnover, whereas α2AR activation inhibits proliferation (Jhaveri et al., 2014). Mechanistically,NE appears to activate a latent stem cell population and then drive progenitor cell division, thus permitting large-scale expansion of hippocampal progenitors (Jhaveri et al., 2010, 2014).

While prior studies have capitalized on pharmacological tools (e.g. neurotoxins, agonists, antagonists) to evaluate the contribution of NE to adult hippocampal neurogenesis, these approaches have some limitations. For example, neurotoxins do not completely lesion the brain noradrenergic system and compensation can occur in surviving cells and afferents, whilst agonists and antagonists target only subsets of ARs and could also exert off-target effects. To determine the consequences of a complete NE ablation on adult hippocampal neurogenesis *in vivo*, we examined knockout mice lacking dopamine β-hydroxylase (*Dbh -/-*), which is required for NE synthesis (Thomas et al., 1995). *Dbh -/-* mice display several phenotypes consistent with impaired adult hippocampal neurogenesis including increased seizure susceptibility, resistance to the behavioral effects of antidepressant drugs, and impairments in some types of learning and memory (Thomas and Palmiter, 1997; Szot et al., 1999; Cryan et al., 2004). As a positive control, we compared our genetic approach with the canonical noradrenergic neurotoxin DSP-4, which we previously showed reduces neurogenesis in rats (Kulkarni et al., 2002).

## 2. Materials and Methods

### 2.1 Animals

Adult male C57BL/6N mice (2-3 months) bred in the TIFR animal colony, and Dbh-/- mice (2-3 months) bred in the Emory University Colony and maintained on a mixed C57Bl6/J and 129SvEv background were group-housed and kept on a 12h light/dark cycle, with access to food and water *ad libitum*. Heterozygous littermates (*Dbh +/-*), which have normal NE content and are phenotypically indistinguishable from wild-type (*Dbh +/+*) mice were used as controls for the *Dbh -/-* mice (Thomas et al., 1995). To prevent embryonic lethality observed in Dbh-/- mice, pregnant Dbh+/-dams were administered the AR agonists, isoproterenol, and phenylephrine (20 μg/ml each) along with vitamin C (2 mg/ml) from embryonic day 9.5 (E9.5)– E14.5, and L-3,4-dihydroxyphenylserine (DOPS; 2 mg/ml + vitamin C; 2 mg/ml) in drinking water from E14.5 until birth. All treatments on C57BL/6N mice were performed at TIFR, while all treatments on *Dbh* mice were performed at Emory University. All experiments were conducted in accordance with the National Institute of Health Guide for the Care and Use of Laboratory Animals and were approved by the TIFR and Emory University Institutional Ethics Committees.

### 2.2 BrdU labeling paradigms and drug treatments

To assess ongoing proliferation of adult hippocampal progenitors, Dbh+/- and Dbh-/- mice (n = 6 per group) received a single intraperitoneal (i.p.) injection of the mitotic marker, bromodeoxyuridine (BrdU; 100 mg/kg; Sigma-Aldrich, USA) and were sacrificed by perfusion 2 h later. To examine the post-mitotic survival of adult hippocampal progenitors, Dbh+/- and Dbh- /- mice (n = 6 per group) were sacrificed four weeks following a single BrdU (100 mg/kg i.p.) administration. To pharmacologically deplete NE levels, the noradrenergic neurotoxin N-(2-Chloroethyl)-N-ethyl-2-bromobenzylamine hydrochloride (DSP4) (10 mg/kg, i.p., Sigma-Aldrich) or vehicle (0.9% saline) was administered to C57BL/6N mice (n = 4 per group) once daily for three days as previously described (Kulkarni et al., 2002). Fluoxetine (5 mg/kg, i.p., kind gift from IPCA laboratories, India) was administered 30 min prior to DSP-4 or vehicle administration to protect serotonergic neurons. Mice received an injection of BrdU (100 mg/kg, i.p.) 3 days after the final DSP-4/vehicle treatment, a time-point associated with effective NE depletion, and were sacrificed 2h after BrdU treatment to assess the influence of pharmacological NE depletion on adult hippocampal progenitor proliferation.

### 2.3 Immunohistochemistry and Immunofluorescence

Mice were sacrificed via transcardial perfusion with 4% paraformaldehyde, brains were dissected and coronal sections (50 μM) were generated on a vibratome (Leica, Germany). BrdU immunohistochemistry was carried out as described previously (Kulkarni et al., 2002). In brief, brain sections were subjected to DNA denaturation (50% formamide/2X Sodium citrate buffer; 2h at 65 °C), followed by acid hydrolysis (2N HCl; 30 mins at 37 °C), blocking 10% horse serum (Gibco, USA) and incubation with rat anti-BrdU antibody (1:500; Axyll Labs, USA). Sections were subjected to washes and incubation with biotinylated goat anti-rat antibody (1:500; Vector Laboratories, USA) for 2h, followed by signal amplification with an Avidin-Biotin complex (Vector Laboratories) and visualization of the signal using diaminobenzidine (Sigma-Aldrich) as a substrate. Sections were mounted using a DPX mountant (Sigma-Aldrich) and viewed using a Zeiss Axioskop 2 plus microscope (Germany). For immunohistochemical and immunofluorescent detection of endogenous markers of adult hippocampal progenitors, polysialylated neural cell adhesion molecule (PSA-NCAM), doublecortin (DCX), Stathmin, TUC-4 (TOAD-64/Ulip/CRMP), brain sections were blocked with 10% horse serum (Gibco) in 0.3% PBTx, followed by overnight incubation with primary antibodies: 1) mouse PSA-NCAM (1:500; a kind gift from Professor T. Seki, Juntendo University School of Medicine, Tokyo, Japan); 2) goat DCX (1:250; Santa Cruz Biotechnology, USA); 3) rabbit Stathmin (1:250; Calbiochem, USA); 4) rabbit TUC-4 (1:250; Chemicon, USA). Following serial washes, sections were incubated with one of the following secondary antibodies: 1) donkey anti-mouse 488 (1:250; Invitrogen, USA); 2) donkey anti-goat 488 (1:250; Invitrogen); 3) donkey anti-rabbit 488 (1:250, Invitrogen). To visualize immunofluorescence, sections were mounted in Vectashield (Vector Labs) and viewed using Zeiss LSM 5 Exciter Confocal microscope (Germany).

### 2.4 Cell counting

All cell counting analyses were performed by an experimenter blind to the treatment conditions. Sections spanning the rostrocaudal extent of the hippocampus (Bregma -1.22 to-3.8mm) were counted using a modified unbiased stereology protocol as described previously (Malberg et al., 2000). In brief, every sixth hippocampal section was processed for BrdU immunohistochemistry (8-10 sections/animal), and the total number of BrdU-positive cells in the subgranular zone (SGZ)/granule cell layer (GCL) was estimated by multiplying the total number of BrdU cells counted by the section periodicity (6). Quantification was performed on the Zeiss Axioskop 2 Plus microscope at a magnification of 400X. Quantitation of PSA-NCAM, DCX, Stathmin, and TUC-4-positive cells within the subgranular zone (SGZ)/granule cell layer (GCL) (4-6 sections/animal) was done using a Zeiss LSM 5 Exciter confocal microscope at a magnification of 400X.

### 2.5 Statistical analysis

Results were subjected to statistical analysis using the unpaired Student’s *t*-test (Prism, GraphPad, USA). Data are expressed as mean ± standard error of the mean (S.E.M) and statistical significance was set at *p* < 0.05.

## 3. Results

### 3.1 Dbh-/- mice do not exhibit altered proliferation, survival, or maturation of adult hippocampal progenitors

Given prior evidence indicating a role for NE in the regulation of adult hippocampal neurogenesis, we sought to address the consequences of a genetic loss of NE via the deletion of the *Dbh* gene. We first assessed ongoing adult hippocampal progenitor proliferation in Dbh+/- and Dbh-/- mice using the mitotic marker, BrdU to label dividing hippocampal progenitors (Fig. 1A). BrdU immunohistochemistry followed by cell counting analysis revealed no change in the number of BrdU-positive progenitor cells within the SGZ in Dbh -/- mice, as compared to Dbh +/- mice (Fig. 1C). We next assessed whether the post-mitotic survival of adult hippocampal progenitors is altered in Dbh-/- mice, by counting the number of surviving BrdU-positive hippocampal progenitors that persist within the GCL of the DG subfield 28 days following a BrdU pulse (Fig. 1B). We observed no difference in the numbers of BrdU-positive progenitors that were observed in the GCL between Dbh -/- mice and controls (Fig. 1D).

**Figure 1:**
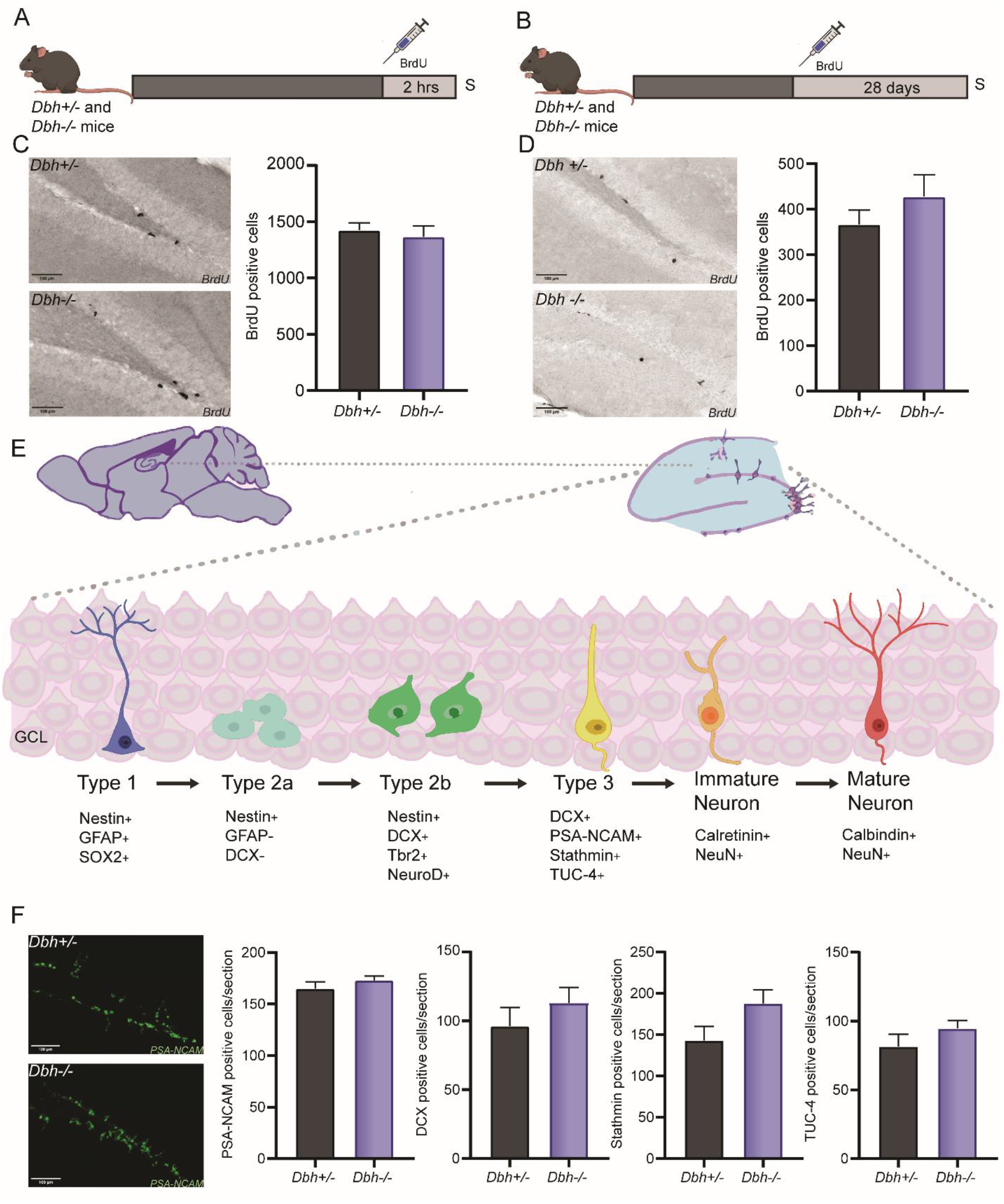
Stage-specific characterisation of hippocampal progenitor markers in DBH +/- and -/- mice. (A) Shown is the schematic representation of the paradigm for labelling hippocampal progenitors with mitotic maker BrdU to assess the changes in hippocampal progenitor proliferation in the dentate gyrus subfield in the hippocampus of Dbh+/- and -/- mice. (B) Shown is the schematic representation of the paradigm for labelling hippocampal progenitors with mitotic maker BrdU to assess the changes in hippocampal progenitor survival in the dentate gyrus subfield in the hippocampus of Dbh+/- and -/- mice. (C) Representative images of BrdU labelling in the dentate gyrus of Dbh+/- and -/- mice for the proliferation paradigm. No changes were seen in the number of BrdU labelled cells between Dbh+/- and -/- mice. (D) Representative images of BrdU labelling in the dentate gyrus of Dbh+/- and -/- mice for the survival paradigm. The number of granule cells labelled with BrdU remained unaltered between Dbh+/- and -/- mice. (E) Shown are the stages of SGZ progenitor development from quiescent neural progenitors to mature granule neurons in the hippocampal along with the expression of stage-specific markers. (F) Representative images of PSA-NCAM positive cells in the dentate gyrus subfield in the hippocampus of Dbh+/- and -/- mice. The number of cells immunopositive for PSA-NCAM remained unchanged between Dbh+/- and -/- mice. Further, no changes were seen in the number of DCX, TUC-4, and Stathmin immunopositive cells between Dbh+/- and -/- mice.

To more extensively characterize the potential impact of genetic loss of NE on the maturation of adult hippocampal progenitors, we assessed the expression of developmental stage-specific markers for adult hippocampal progenitors in Dbh+/- and Dbh-/- mice (Fig. 1E). We performed immunofluorescence studies for PSA-NCAM and DCX, which are reported to first appear in the type 2b class of hippocampal progenitors and are expressed in the predominantly post-mitotic type 3 class of progenitors and immature granule cell neurons (Bond et al., 2015). The number of PSA-NCAM and DCX-immunopositive cells per section within the SGZ/GCL did not differ between Dbh+/- and Dbh-/- mice (Fig. 1F). We also observed no effect of *Dbh* genotype on the number of TUC-4 positive cells in the SGZ/GCL (Fig. 1F), which is considered to be a marker for late mitotic neuronal progenitors and early postmitotic neurons and has been reported to overlap with PSA-NCAM and DCX in the time-window of expression during stage-specific progression of adult hippocampal progenitor cells (Bond et al., 2015). We also noted no change in stathmin-positive hippocampal progenitor numbers in Dbh-/- mice (Fig. 1F). Collectively, these results reveal that genetic loss of NE does not appear to influence adult hippocampal progenitor turnover, survival, or maturation.

### 3.2 DSP-4 evoked depletion of NE impairs adult hippocampal neurogenesis in mice

Prior evidence indicates that pharmacological approaches to deplete NE, via administration of the noradrenergic neurotoxin DSP-4, evoked a significant decline in adult hippocampal progenitor cell division in rats (Kulkarni et al., 2002). The failure of *Dbh* knockout to affect hippocampal neurogenesis in mice suggested that either NE is important for neurogenesis in rats but not mice, or that neurotoxic lesions reveal a critical role for the noradrenergic system in a way that genetic manipulations do not. Administration of DSP-4 to wild-type C57BL6/N mice (Fig. 2A) resulted in a significant reduction in the number of BrdU-positive proliferating hippocampal progenitors in the SGZ/GCL (Fig. 2B). Despite the decline in adult hippocampal progenitor proliferation in mice following DSP-4 treatment, we observed no significant difference in DCX (Fig. 2C) or PSA-NCAM-positive (Fig. 2D) adult hippocampal progenitor numbers in the SGZ/GCL at the time-point examined.

**Figure 2:**
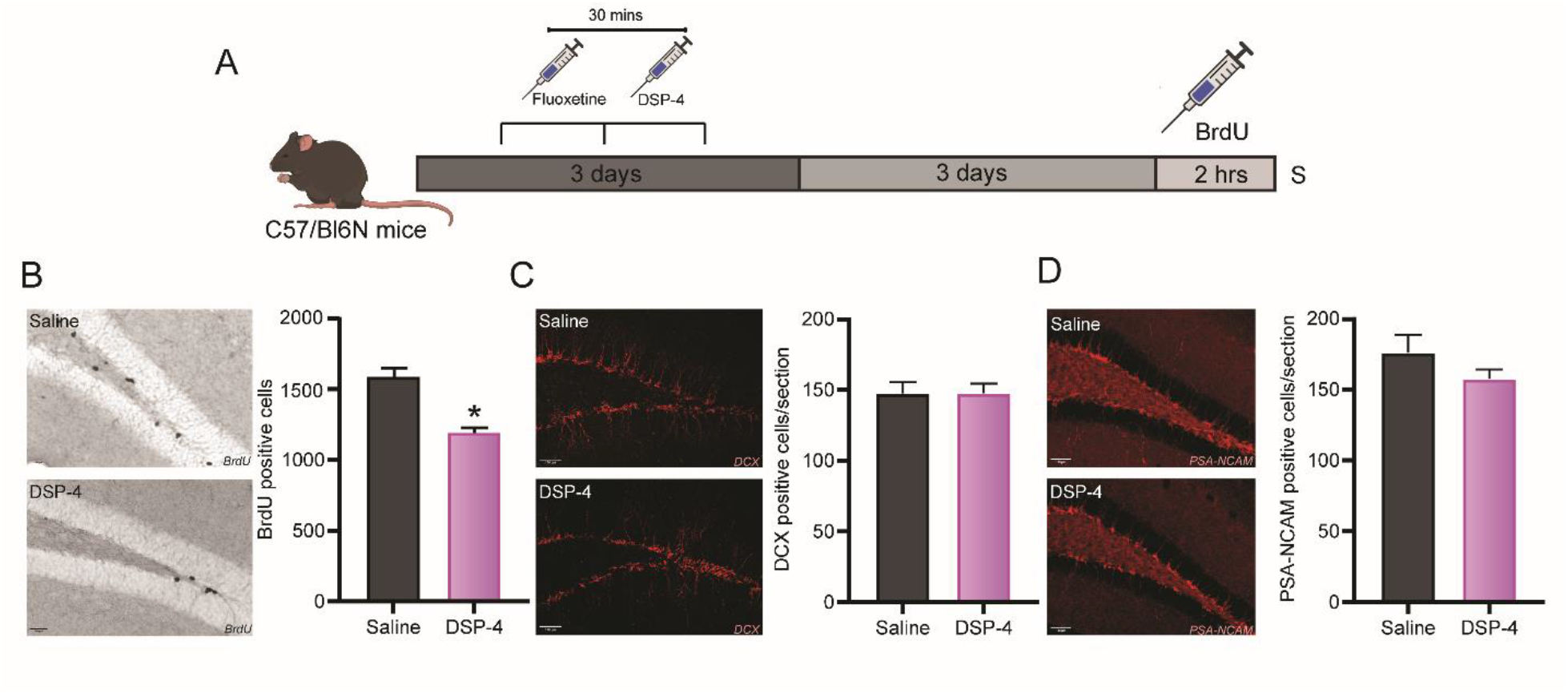
Stage-specific characterisation of hippocampal progenitor markers in C57BL6N mice treated with 10mg/kg DSP-4. (A) Shown is the schematic representation of the paradigm for labelling hippocampal progenitors with mitotic maker BrdU to assess the changes in hippocampal progenitor proliferation in the dentate gyrus subfield in the hippocampus of vehicle or DSP-4 treated mice. (B) Representative images of BrdU labelling in the dentate gyrus of C57BL6N mice treated with vehicle or DSP-4. The number of BrdU labelled cells was significantly reduced in the vehicle or DSP-4 treated mice. (C) Representative images of DCX positive cells in the dentate gyrus of C57BL6N mice treated with vehicle or DSP-4. No changes were seen in the number of DCX immunopositive cells between vehicle or DSP-4 treated mice. (D) Representative images of PSA-NCAM positive cells in the dentate gyrus of C57BL6N mice treated with vehicle or DSP-4. The number of cells immunopositive for PSA-NCAM remained unchanged between vehicle or DSP-4 treated mice.

## 4. Discussion

Substantial evidence has uncovered an important role for NE in the modulation of adult neurogenesis, in particular in the hippocampal neurogenic niche. NE can influence distinct stages of development of adult hippocampal progenitors, with robust effects on latent stem cell activation that can enhance the available progenitor pool (Jhaveri et al., 2010; Yanpallewar et al., 2010; Jhaveri et al., 2014). Further, NE is reported to enhance adult hippocampal progenitor cell division, and increase neuronal differentiation (Jhaveri et al., 2010, 2014). Pharmacological lesioning of LC noradrenergic afferents is also associated with a robust decline in adult hippocampal progenitor proliferation (Kulkarni et al., 2002; Coradazzi et al., 2016). Counter to prior data using pharmacological approaches and our expectations, *Dbh -/-* mice that exhibit a complete genetic loss of NE did not show any alteration in ongoing adult hippocampal neurogenesis. These results were rather surprising given that NE is implicated in multiple stages of adult progenitor developmental progression in the hippocampal niche including stem cell activation, progenitor turnover, and cell fate specification. In contrast, lesions of the LC by the selective neurotoxin DSP-4 significantly reduced adult hippocampal progenitor proliferation in mice, similar to our prior results in rats (Kulkarni et al., 2002).

Our results reveal that the number of proliferating hippocampal progenitors labeled with a pulse of the mitotic marker BrdU is unaltered both at the short time-point of 2h and also the longer duration time-point of 28 days, post the BrdU pulse in *Dbh -/-* mice. This demonstrates that both hippocampal progenitor cell division and the post-mitotic survival of labeled hippocampal progenitors are not different in animals with a genetic loss of NE. Quantitative analysis also revealed that the numbers of PSA-NCAM, DCX, stathmin, and TUC-4 positive cells, which are expressed by type 2b and type 3 neuroblasts in the hippocampal neurogenic niche (Kempermann et al., 2004), were also not changed in *Dbh -/-* mice, as compared to *Dbh +/-* mice. We also did not observe any overt differences in newborn neuron morphological maturation as revealed by the dendritic complexity of DCX-positive immature neurons in *Dbh -/-* mice. Our control group for all our studies was *Dbh +/-* mice that exhibit NE levels that do not differ from the wild-type (Thomas et al., 1995).

We have previously shown that pharmacological depletion of NE by lesioning noradrenergic neurons with DSP-4 in rats results in a marked reduction in adult hippocampal progenitor proliferation (Kulkarni et al., 2002). This finding has thus far not been replicated in mice, raising the possibility of species differences in the neurogenic decline noted following NE depletion. We carried out studies with DSP-4 administration in mice and noted a significant decrease in progenitor turnover, indicating that lesioning of noradrenergic afferents exerts similar effects in both rats and mice. This clarifies that the absence of effects on hippocampal neurogenesis in *Dbh -/-* mice are not simply a consequence of species differences in the neurogenic effects of NE. This conclusion is further bolstered by prior evidence from studies that indicate robust neurogenic effects of NE elevation and stimulation of β3ARs on adult hippocampal progenitors both *in vivo* and *in vitro* in mice (Jhaveri et al., 2010, 2014). Given these prior findings, and our current results that demonstrate a significant effect of pharmacological depletion of NE on hippocampal neurogenesis in mice, it is intriguing that the *Dbh -/-* mice, which have a completely lack NE, continue to exhibit normal ongoing adult hippocampal neurogenesis.

There are several important differences between genetic and neurotoxic approaches to NE depletion that could account for the discrepancies in the results we obtained from the two approaches. First, *Dbh -/-* mice lack NE from birth, and compensations during postnatal development may occur that do not arise in the DSP-4 model, which involves lesioning in adulthood and testing soon after. Second, because DBH converts dopamine to NE, *Dbh -/-* mice synthesize dopamine instead of NE in their noradrenergic neurons (Weinshenker et al., 2002). Much of the dopamine in the hippocampus is derived from LC neurons even in normal mice, and dopamine has been implicated in modulating adult hippocampal neurogenesis, with studies reporting a decline in progenitor proliferation in animals with the pharmacological lesion of dopaminergic projections (Höglinger et al., 2004). Thus, one speculative possibility is that the excess dopamine may substitute for NE in the context of adult hippocampal neurogenesis in *Dbh-/-* mice. Third, LC neurons contain many neuropeptides besides NE with neurotrophic properties, including brain-derived neurotrophic factor (BDNF), galanin, and neuropeptide Y (Poe et al., 2020). LC neuropeptide levels are spared in *Dbh-/-* mice (Poe et al., 2020), while neurotoxin LC lesions deplete both NE and other LC-derived neuromodulators. The loss of these other neurotrophic molecules may impact adult hippocampal neurogenesis in ways that specific NE depletion cannot. Finally, DSP-4 lesions involve the physical deterioration of LC fibers and terminals and neuroinflammation, while *Dbh -/-* mice have completely intact and healthy LC neurons and projections. Local degeneration of LC fibers in the hippocampus and associated neuroinflammatory processes could impair ongoing neurogenesis in a NE-independent manner (Monje et al., 2003). It is also of interest to note that we have previously shown that *Dbh-/-* mice exhibit changes in plasticity-associated markers in other neurogenic niches (Vadodaria et al., 2017). For example, expression of the plasticity markers DCX and PSA-NCAM was found to be significantly elevated in the piriform cortex of *Dbh-/-* mice, indicating that NE and potential compensatory mechanisms in its absence may not function equivalently across all brain regions (Vadodaria et al., 2017). *Dbh-/-* mice exhibit diverse behavioral changes spanning from altered seizure susceptibility to motor dysfunction, including a treatment resistance phenotype to pharmacological antidepressants (Thomas and Palmiter, 1997; Szot et al., 1999; Cryan et al., 2004). Further, they also exhibit changes in ARs, with reports of a decline in the α2AR in limbic brain regions including the hippocampus, and an increase in βAR binding(Sanders et al., 2006). This is particularly intriguing given that the α2AR is known to inhibit adult hippocampal progenitor proliferation, whilst the β3AR stimulates latent stem cell activation and progenitor turnover, raising the possibility of the tipping of balance towards a stimulatory drive on adult hippocampal progenitor turnover. The absence of neurogenic effects noted in *Dbh-/-* mice could arise due to a variety of reasons, and future experiments will be required to precisely pinpoint the underlying mechanisms.

## Acknowledgments

This work was supported by intramural funding from the Tata Institute of Fundamental Research and the Department of Atomic Energy, Mumbai (RTI4003 to VV) and the National Institute of Health (AG061175 and AG079199 to DW). We thank Dr. Shital Suryavanshi, K.V. Boby, and the animal house staff at the Tata Institute of Fundamental Research, Mumbai, for technical assistance.

## Author contributions

DK, KCV, KSR, YEO, LCL, KAFT, SSS, BFAH contributed to animal experiments and data analysis. DW and VV designed all experiments and contributed to data analysis. DK, DW and VV wrote the manuscript.

